# Assessment of atrioventricular nodal reentrant tachycardia inducibility with alternative pacing methods: Computer simulation

**DOI:** 10.1101/2025.11.27.690941

**Authors:** Maxim Ryzhii, Elena Ryzhii

## Abstract

**Background and objective:** In clinical and experimental electrophysiology, the standard S1S2 stimulation protocol is widely used to assess the inducibility of reciprocal tachycardias. However, this method may lead to over- or under-estimation of atrioventricular nodal reentrant tachycardia (AVNRT) susceptibility. Computational modeling enables a detailed examination of these limitations and the design of more physiologically accurate stimulation approaches. This study evaluates alternative stimulation methods using a compact, multifunctional rabbit AV node model with autonomic nervous system (ANS) control.

**Methods:** The model, based on experimental data, incorporates dual (fast and slow) pathway conduction and simulates the propagation of nodal electrical excitation. Alternative stimulation protocols were implemented and compared with the standard S1S2 approach at both atrial and His bundle stimulation sites.

**Results:** Simulations yielded AV node ladder diagrams, effective refractory periods, and echo response zones across the full range of ANS modulation. The proposed methods enabled accurate characterization of nodal pathway refractory periods during retrograde conduction and identified physiological conditions leading to intranodal reentry. Compared with the S1S2 protocol, the alternative approaches reduced misestimation of AVNRT inducibility and provided clearer separation between echo beats and sustained reentry.

**Conclusions:** The proposed stimulation methods, particularly during His bundle pacing, offer a more accurate and physiologically grounded assessment of AVNRT inducibility than the conventional S1S2 protocol, improving the understanding of nodal conduction dynamics and arrhythmia mechanisms.

## 1. Introduction

The atrioventricular (AV) node is a key component of the cardiac conduction system, located in the interatrial septum near the coronary sinus. Acting as the electrical gateway between the atria and ventricles, it ensures coordinated contraction and efficient blood circulation [1]. Electrical impulses from the sinus node (SN) travel through the AV node to the His bundle, which transmits signals to the Purkinje fibers and ultimately the ventricular myocardium. A physiological delay introduced by the AV node allows sufficient time for atrial contraction and ventricular filling before ventricular depolarization occurs. In addition to its conduction role, the AV node acts as a rate regulator, filtering high-frequency impulses during atrial arrhythmias such as atrial fibrillation and flutter. When the sinus node fails, the AV node can serve as a secondary pacemaker to preserve cardiac output. AV nodal dysfunction can lead to conduction blocks resulting in bradycardia, and in severe cases, may necessitate pacemaker implantation.

Structurally, the AV node is composed of slow-conducting nodal cells that differ from the fast-conducting fibers of the atria and ventricles. It is innervated by the autonomic nervous system (ANS), which modulates conduction velocity and refractory periods through sympathetic and parasympathetic input [2]. Functionally, the AV node exhibits dual-pathway conduction [3, 4], consisting of a fast pathway (FP) and at least one slow pathway (SP). The FP conducts impulses more rapidly but has a longer effective refractory period (ERP), whereas the SP conducts impulses more slowly but recovers more quickly. This dual-pathway architecture plays a protective role but also underlies the mechanism of AV nodal reentrant tachycardia (AVNRT) [5, 6, 7].

Slow-fast (typical) AVNRT occurs when a premature atrial impulse encounters refractory FP and is redirected anterogradely through SP. Once the FP recovers, the impulse may re-enter it retrogradely, creating a reentrant loop that leads to rapid, repetitive activation. In contrast, fast-slow (atypical) AVNRT is usually caused by a premature His bundle stimulus and retrograde conduction through the SP [8, 9].

The timing of the premature impulse, along with the prevailing autonomic tone, significantly influences the initiation, maintenance, and termination of AVNRT [10, 11]. This condition is most often induced during periods of heightened sympathetic activity and may terminate spontaneously or through increased vagal tone [11, 12], which can be achieved either pharmacologically or through vagal maneuvers.

The difference in anterograde and retrograde ERPs between the FP and SP, in other words, the induction window formed by the activity of the ANS, plays a central role in the occurrence of reentry and the generation of echo rhythms [12, 13].

The properties of anterograde conduction and ERPs in the AV node have been extensively studied in various contexts, including animal experiments [14, 15, 16], patient studies [17, 18], and through computational approaches. These computational methods encompass detailed ionic models [19, 20], simplified cell models [11, 21], functional mathematical models [14, 22], and network models [23, 24]. However, the retrograde properties of the AV node have received comparatively less attention [13, 16, 25, 26].

One of the most widely used experimental tools in electrophysiological studies is the S1S2 pacing protocol. This protocol involves delivering a series of 8 to 10 regular stimuli (S1) at a fixed cycle length of 300 to 600 ms, followed by a premature extrastimulus (S2). The interval for the test S2 stimulus is progressively shortened until conduction fails, allowing for the identification of the ERP of the tissue. The S1S2 pacing protocol is crucial for examining the dual-pathway physiology of the AV node, assessing conduction delays, and evaluating both effective and functional refractory periods. In both clinical and experimental contexts, S1S2 and S1S2S3 pacing protocols are commonly utilized to investigate the inducibility of reentrant tachycardias, such as AVNRT and atrioventricular reentrant tachycardia. These protocols also help differentiate the anatomical locations of reentrant circuits [5, 27, 28].

Because AV nodal ERP measurements are sensitive to the choice of pacing protocol, different stimulation techniques used during electrophysiological studies may alter the apparent conduction properties, potentially leading to either exaggeration or depreciation of the AVNRT onset possibility.

Several pacing methods have been proposed to improve the accuracy of assessing the occurrence of supraventricular reciprocating tachyarrhythmias (see, for example, [26, 29, 30, 31]). Since AV nodal ERP measurements are sensitive to the pacing protocol used, different stimulation techniques during electrophysiological studies can affect the apparent conduction properties. This variability may either exaggerate or underestimate the likelihood of AVNRT onset [32].

Therefore, interpreting nodal conduction curves, which are used to characterize AV nodal function across different species and conditions, requires careful consideration of pacing methodology. This is particularly important when evaluating the effects of autonomic modulation or pharmacological agents.

Pacing from the atrial side generally allows for stable overdrive of sinus node activity, minimizing its influence on AV nodal measurements. This approach is suitable for accurately assessing anterograde conduction and ERP in both unmodified and ablated AV nodes. In contrast, results from His bundle (para-Hisian) pacing may be affected by the anterograde conduction initiated in the sinus node or atria. As we demonstrate below, the retrograde His-atrial and atrial-atrial conduction intervals can vary significantly depending on the stimulation method used. This variation can lead to differences in the assessment of the probability of AVNRT occurrence.

Recent work from our group introduced a computational model of the rabbit AV node that incorporates dual-pathway physiology controlled by the ANS, along with visualizations of Lewis ladder diagrams [11]. In that study, we modeled the modulation of refractoriness using a single coefficient that affects conduction delay and intrinsic pacemaker activity. We demonstrated how autonomic tone impacts the dynamics of slow-fast and fast-slow AVNRT, highlighting key conditions that either promote or inhibit reentry. However, a detailed analysis of anterograde and retrograde ERPs in the FP and SP, as well as their relationship to the inducibility of AVNRT, was not explored.

In the present study, we aim to investigate how specific ERP measurements affect the assessment of AVNRT induction using our AV node model. We propose alternative two-stage pacing methods applied from both the atrial side and the His-bundle side of the AV node, and compare these methods with the S1S2 standard pacing protocol. Our goal is to identify the limitations of current standard stimulation techniques that may lead to misinterpretation of AVNRT inducibility, either by unintentionally favoring reentry or by suppressing it through premature pathway block. This work seeks to provide electrophysiologists with valuable insights into the effectiveness and reliability of both standard and alternative pacing protocols. Ultimately, we aim to enable a more accurate assessment of AV nodal conduction and the detection of AVNRT inducibility.

## 2. Model and Methods

### 2.1. AV Node Structure and Cardiac Cell Model

The AV node and surrounding conduction system (see Fig. 1) were modeled using the Aliev–Panfilov excitable cell model [33], adapted for autonomic nervous system (ANS) modulation following [11, 34]. Each model cell is described by the following reaction–diffusion system of differential equations:

**Figure 1:**
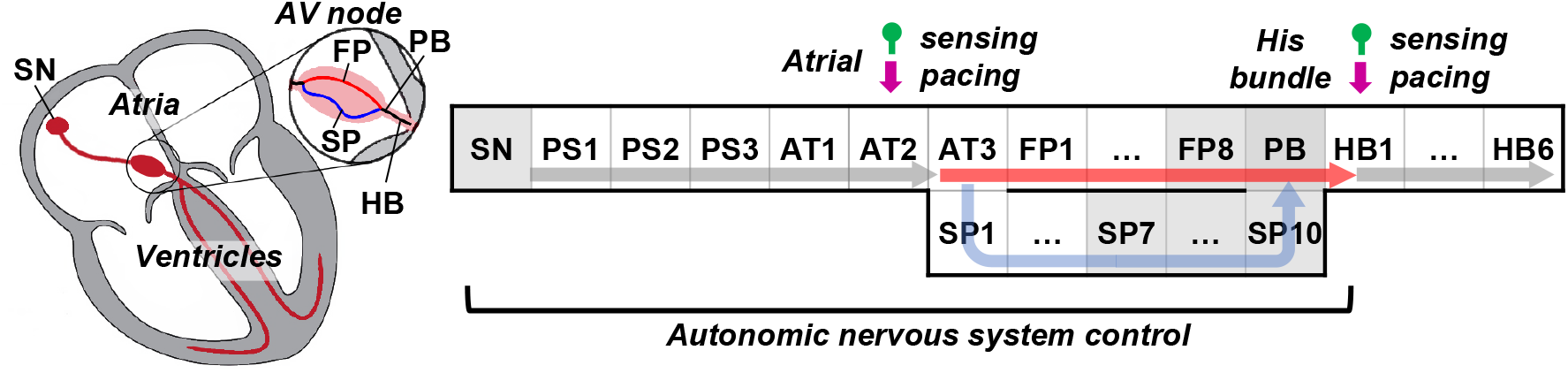
Schematic of the cardiac conduction system and the rabbit atrioventricular node model. SN indicates the sinus node, PS the peripheral sinus node cells, AT the atrial cells, FP the fast pathway, SP the slow pathway, PB the penetrating bundle, and HB the His bundle. Green pin markers indicate sensing sites (AT2 for atrial pacing and HB1 for His bundle pacing), and vertical magenta arrows indicate stimulus application. Arrows within the structure show normal anterograde conduction through the fast pathway. Pacemaker cells are shown in gray.

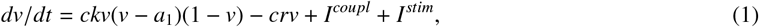

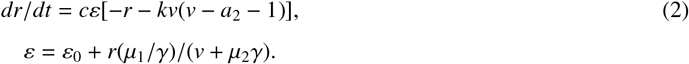

Here, 0 < *v* < 1 represents the dimensionless transmembrane potential, and *r* > 0 is the recovery gating variable. The time-scaling factor *c* sets physical time, while *k* controls the magnitude of the transmembrane ionic current. The depolarization and repolarization currents are represented by the first and second terms in Eq. (1). The term *I*^*coupl*^ describes intercellular diffusion currents, including coupling asymmetry [21], and *I*^*stim*^ denotes externally applied stimuli delivered to atrial or His bundle sites (Fig. 1). Parameters *ε*_0_, *a*_1_, *a*_2_, *µ*_1_, and *µ*_2_ jointly determine conduction velocity, excitability, and refractory behavior. Positive *a*_1_ sets the excitation threshold, while negative *a*_1_ yields intrinsic pacemaker activity, as in sinus and junctional cells (Fig. 1) [35].

### 2.2. Autonomic Nervous System Modulation

ANS effects on conduction and refractoriness are modeled through the parameter *γ* [11, 34]:

- *γ* = 1.7 corresponds to balanced autonomic tone and normal rhythm,
- *γ <* 1.7 represents increased sympathetic tone, producing shorter action potential durations, shortened effective refractory periods (ERPs), reduced nodal conduction times, and accelerated intrinsic sinus and junctional rates [13],
- *γ* > 1.7 corresponds to vagal dominance, generating opposite effects.

The same *γ* value is applied to all cells from the sinus node to the His–Purkinje system, including junctional pacemaker regions [4].

### 2.3. AV Node Model Variants

We used three variants of the AV node model described in [11]. All variants share similar anterograde dual-pathway properties, with the fast pathway (FP) exhibiting a longer ERP than the slow pathway (SP). The variants differ in their retrograde conduction characteristics:

- Variant 1: With dominant sympathetic tone, the retrograde ERP of the FP is equal to that of the SP. As *γ* increases, the FP refractoriness progressively lengthens.
- Variant 2: The FP maintains a consistently longer retrograde ERP than the SP across the full range of *γ*.
- Variant 3: At elevated sympathetic tone, the SP exhibits a longer retrograde ERP. When *γ >* 1.5, this relationship reverses, producing an inverse ERP dependence.

These differences play a crucial role in predicting retrograde propagation, echo beats, and conditions favoring initiation of AV nodal reentrant tachicardia.

### 2.4. Pacing Protocols and ERP Measurement

#### 2.4.1. S1S2 Pacing for ERP Determination

The standard S1S2 protocol consists of nine S1 stimuli delivered at a cycle length slightly shorter than the spontaneous sinus interval determined by the current ANS state. A premature S2 stimulus follows, with the S1–S2 interval shortened in 1 ms steps. AV nodal conduction time is measured between the AT2 and HB1 cells (Fig. 1). Stimuli are 1 ms (atrial) or 2 ms (His bundle) in duration, with amplitudes set to 1.3× threshold. The ERP of a pathway is defined as the longest S1–S2 interval that fails to propagate between the proximal and distal ends of the AV node in the relevant direction.

#### 2.4.2. Atrial Pacing Methods

We implemented the following atrial pacing methods:

- A1A2 (standard atrial S1S2) - atrial stimuli overdrive sinus activity and define the A1–A2 coupling interval [Fig. 2(a)], and
- SbA2 (sinus-based atrial S2) - similar to A1A2, except the A1–A2 interval is measured relative to the arrival of the sinus beat (Sb) at the proximal AV node (AT2 cell) [Fig. 2(b)].

**Figure 2:**
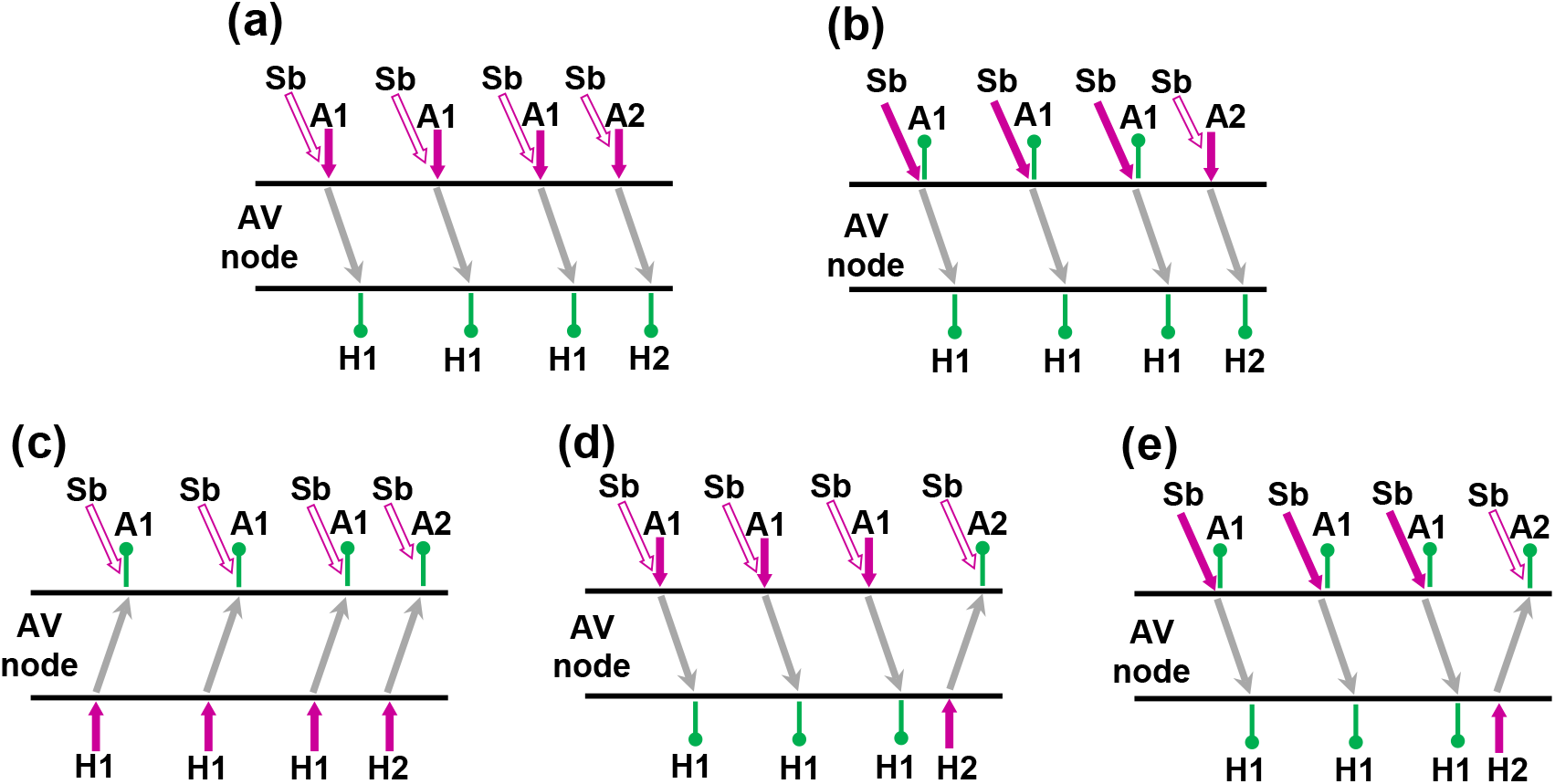
Pacing schemes. (a) A1A2, standard atrial pacing; (b) SbA2, sinus beats (Sb) followed by an A2 test stimulus; (c) H1H2, standard His bundle pacing; (d) A1H2; and (e) SbH2, sinus beats followed by an H2 test stimulus. Filled arrows indicate stimulation, opaque arrows represent overdriven sinus beats, and pinpoints mark sensing sites.

#### 2.4.3. His Bundle Pacing Methods

We examined three His bundle stimulation protocols:

- H1H2 (standard His bundle S1S2) - basic and test stimuli are applied at HB1 cell [Fig. 2(c)].
- A1H2 (atrial S1, His bundle S2) - basic A1 stimuli are delivered to AT2 to overdrive sinus rhythm, followed by a test His bundle stimulus H2 at HB1 cell [Fig. 2(d)]. The H1–H2 interval is measured from the arrival of the final basic beat at HB1.
- SbH2 (sinus-based His bundle S2) - sinus activity provides basic anterograde stimuli, and the test H2 stimulus applied similarly to the A1H2 method [Fig. 2(e)].

These alternative protocols allow for more accurate assessment of retrograde conduction and onset of echo beats and AVNRT.

### 2.5. Visualization of Ladder Diagrams

For visualization of the ladder diagrams, we used our original algorithm with the exact activation timing of each model cell. It allows tracing the wavefront propagation for each excitation sequence and in each pathway separately based on the activation time in each cell. The activation times in each model cell were recorded at the level of 0.25 during the action potential upstroke.

### 2.6. Quantitative Assessment of the Proposed Methods

For the proposed His bundle pacing methods we evaluated their effectiveness in the reentrant activity inducibility (identification) compared with standard H1H2 protocol. The over-, underestimation, and total errors of the latter were calculated as

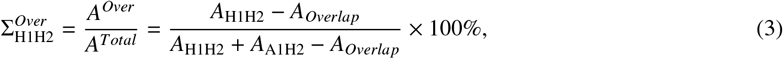

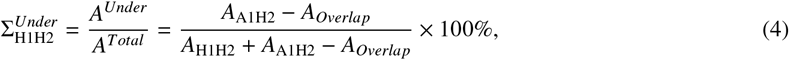

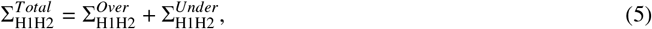

where *A*_H1H2_ and *A*_A1H2_ are the areas on the reentrant activity maps (test pacing interval versus *γ*), produced by the standard H1H2 and new A1H2 (SbH2) methods, respectively, and *A*_*Overlap*_ is the common area for both methods.

### 2.7. Numerical Implementation

Equations (1)–(3) were solved in MATLAB (R2024b, MathWorks Inc., Natick, MA, USA) using the ODE23 solver with parameter values consistent with [11]. The implementation preserves conduction asymmetry across path- ways and allows systematic variation of *γ* to simulate ANS effects.

## 3. Results

### 3.1. Atrial Pacing

#### 3.1.1. Comparison of A1A2 and SbA2 Pacing in Isolated Pathways

Figure 3 demonstrates the comparison of ladder diagrams obtained with the atrial A1A2 and sinus-based atrial SbA2 protocols for a test interval of 240 ms. These diagrams were generated for isolated fast pathway (FP) and slow pathway (SP) configurations by ablating the opposite pathway. In both pacing approaches, the transformation of the A1–A2 test interval into the resulting H1–H2 interval is nearly identical, as emphasized by the dashed arrows. This confirms that the two atrial pacing strategies impose essentially the same anterograde loading conditions on the AV node.

**Figure 3:**
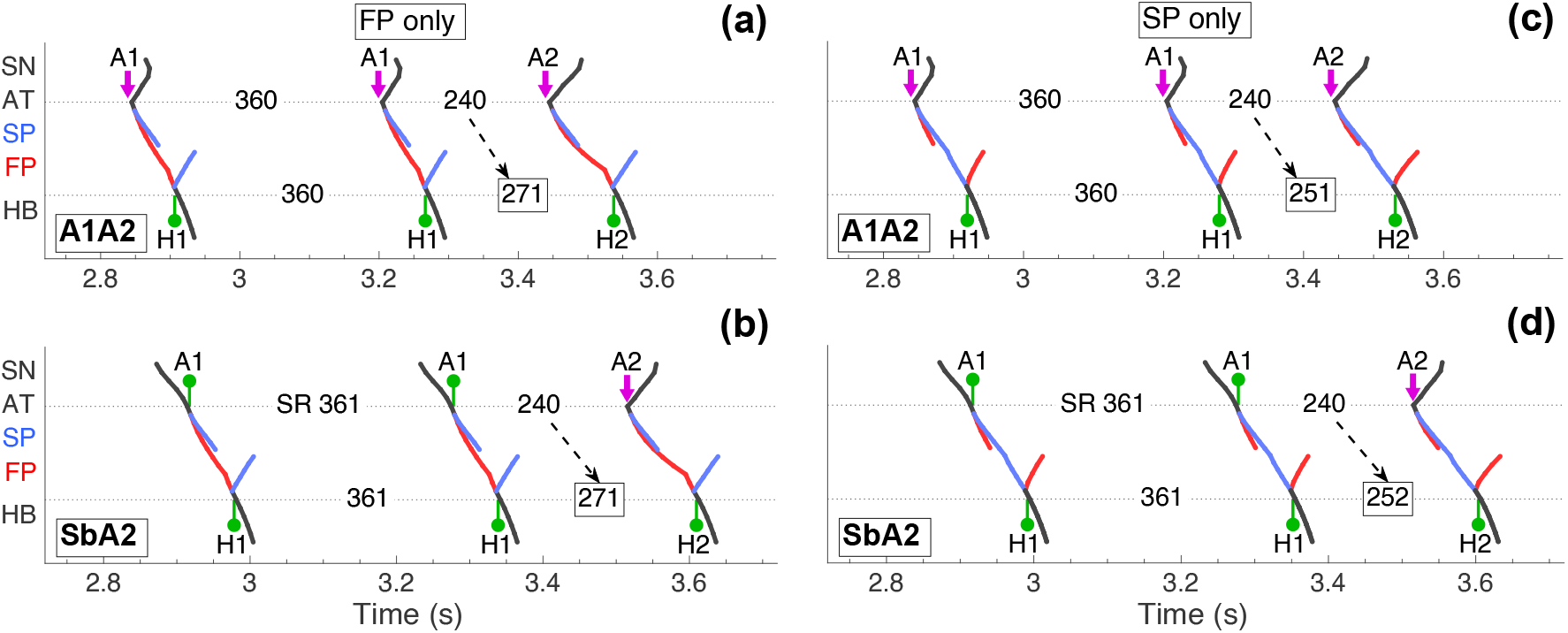
Ladder diagrams for the first model variant obtained using atrial pacing A1A2 and SbA2 pacing methods at autonomic system coefficient *γ* = 1.7 and an H1-H2 test interval of 240 ms. Panels (a) and (b) show conduction with the fast pathway (FP) only after slow pathway (SP) ablation, and panels (c) and (d) show conduction with the SP only after FP ablation. Red tracings correspond to FP activity and blue - to SP activity. Pacing and conduction intervals, including the sinus rhythm (SR) interval in the bottom panels, are given in ms. Magenta arrowheads mark stimulation and green pin markers indicate excitation sensing. Dashed arrows highlight the differences between the imposed A–A test interval and the resulting H–H interval.

#### 3.1.2. Anterograde ERPs Across Autonomic States

The dependence of FP and SP anterograde effective refractory periods (ERPs) on autonomic state, expressed through the coefficient *γ*, and the reentrant activity maps are presented in Fig. 4 for all three model variants. ERPs obtained with A1A2 pacing closely reproduce previously reported results and closely match those of the SbA2 method.

**Figure 4:**
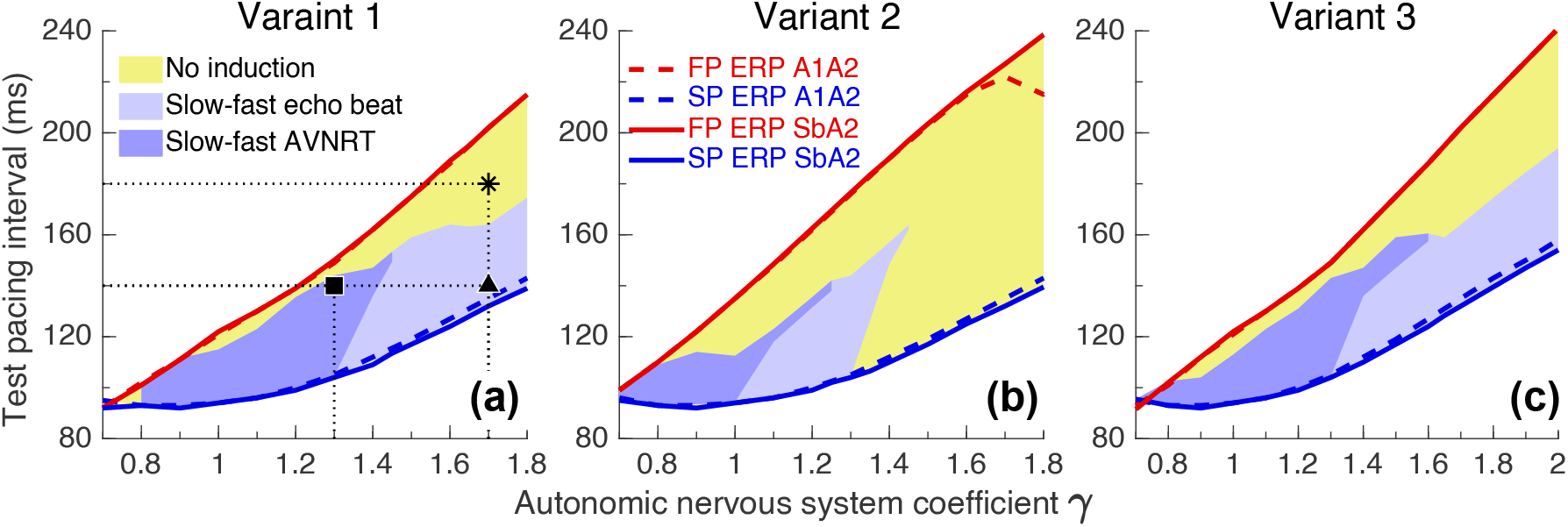
Anterograde effective refractory periods (ERPs) as a function of the autonomic nervous system coefficient *γ*, and reentrant activity maps for the first (a), second (b), and third (c) model variants computed using the standard A1A2 protocol (dashed lines) and the SbA2 protocol (solid lines). Red and blue curves indicate the ERPs of the fast (FP) and slow (SP) pathways. Light and dark blue shading mark zones of slow-fast echo beats and AVNRT induction, and yellow shading identifies non-inducible regions. Square, star, and triangle markers denote the parameter points analyzed in detail in Fig. 5.

Minor discrepancies arise from differences in the coupling current reaching the AT2 cell, which originates from the sinus node in SbA2 pacing but is externally injected during A1A2 pacing.

Regions associated with slow–fast atrioventricular nodal reentrant tachycardia (AVNRT) and echo beats, marked by dark and light shading, appear in similar *γ* ranges for both methods. In areas below the curves representing shorter ERPs, both pathways are in a non-conductive state, whereas in regions above the curves corresponding to longer ERPs, both pathways are conductive. Although the difference in ERP between pathways increases as *γ* increases, the actual regions supporting AVNRT or echo rhythms do not expand proportionally as expected and may even narrow with further increases in *γ*.

#### 3.1.3. Conduction Patterns in the Unmodified AV Node

Ladder diagrams for selected points on the reentrant activity maps of the first model variant are shown in Fig. 5. These examples cover conditions that induce slow–fast AVNRT, generate isolated echo beats, or fail to initiate any reentry. The A1A2 and SbA2 conduction patterns remain nearly identical, except for slight differences in retrograde propagation toward the sinus node. When sympathetic tone is strongly dominant (*γ* ≤ 1.4), with 140 ms test interval FP cells recover by the time the anterograde wavefront in SP reaches the lower pathway segments, enabling successful retrograde propagation through FP and allowing AVNRT to develop [Fig. 5(a,b)]. As parasympathetic activity increases (for example, at *γ* = 1.7), the same 140 ms test interval leads to a complete block of anterograde FP conduction at proximal part of AV node and to single echo beat via FP as shown in Fig. 5(c, d). On the other hand, at a longer test interval of 180 ms and same *γ*, a certain number of FP cells gradually recover from their refractory state, allowing concealed anterograde conduction to emerge in the proximal part of FP [marked by dashed circles in Fig. 5(e,f)], inhibiting retrograde FP conduction somewhere along its way, preventing the onset of echo beats or AVNRT. This effect creates no-induction regions (marked yellow in Fig. 4) adjacent to FP ERP curves on the re-entrant activity maps in all three model variants.

**Figure 5:**
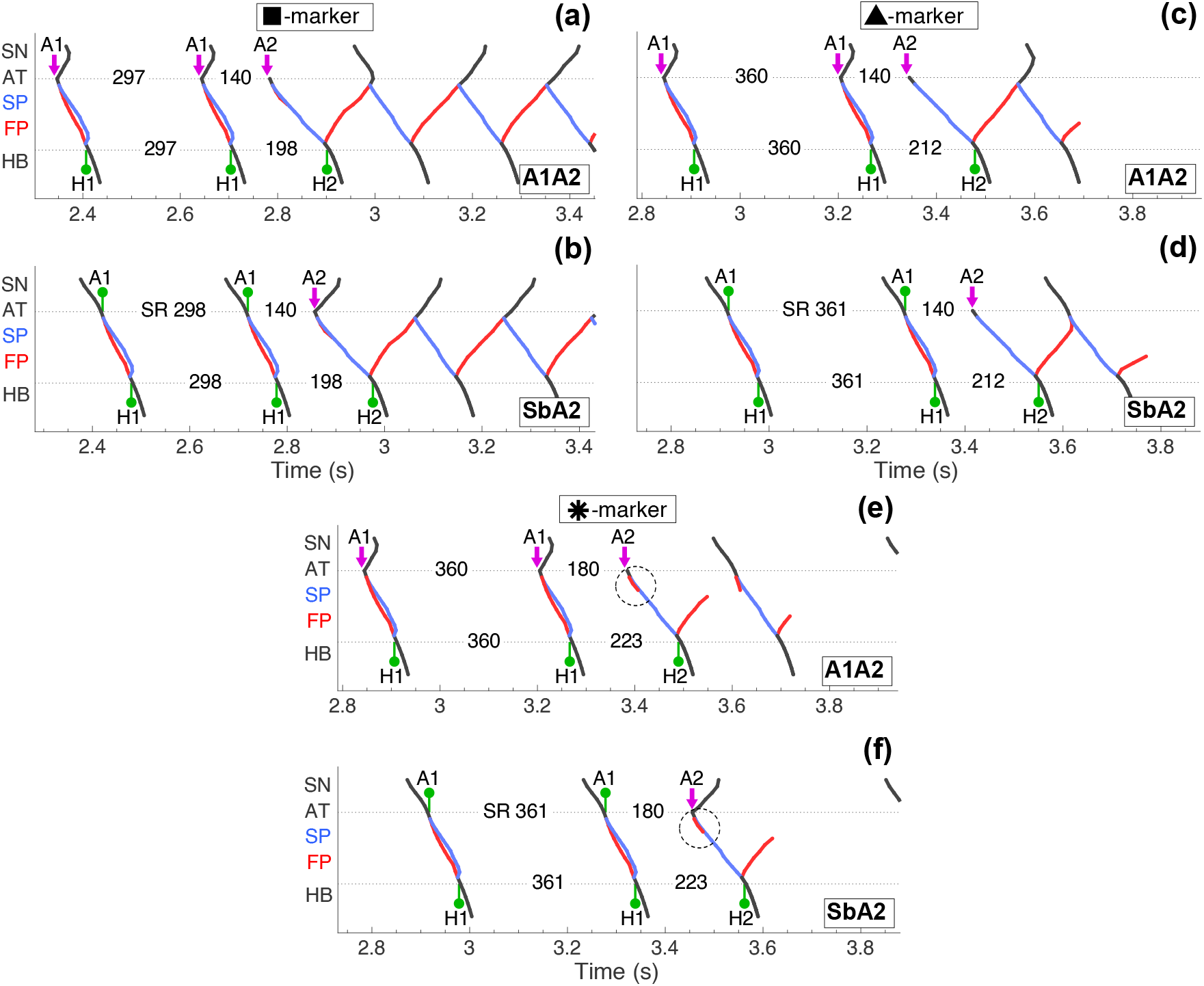
Ladder diagrams for the unmodified AV node in the first model variant during A1A2 and SbA2 atrial pacing. Panels (a) and (b) show slow-fast AVNRT onset at *γ* = 1.3 with a 140 ms test interval [square marker in Fig. 4(a)]. Panels (c) and (d) show echo beats at *γ* = 1.7 and a 140 ms interval (triangle marker). Panels (e) and (f) show absent reentry and no echo beats at *γ* = 1.7 with a 180 ms interval (asterisk marker).

### 3.2. His Bundle Pacing

#### 3.2.1. Comparison of Standard H1H2 and Alternative HIS Bundle Pacing Protocols

Figure 6 provides representative ladder diagrams for His bundle pacing at *γ* = 1.7 and a test interval of 240 ms. For both FP-only and SP-only configurations with ablated opposite pathways, the standard H1H2 protocol produces results that differ markedly from those obtained with the A1H2 and SbH2 methods. In H1H2 pacing, all applied stimuli propagate retrogradely, whereas the alternative protocols impose an anterograde–retrograde transition: basic excitation arrives anterogradely from the atria, while only the test stimulus travels retrogradely. This shift extends the interval between consecutive activations of the same pathway segment, enhancing recovery from refractoriness and thereby shortening the measured ERPs for both pathways. The transformation of the H1–H2 interval into A1–A2 interval, highlighted in the diagrams, demonstrates how retrograde recovery is facilitated under A1H2 and SbH2 pacing.

**Figure 6:**
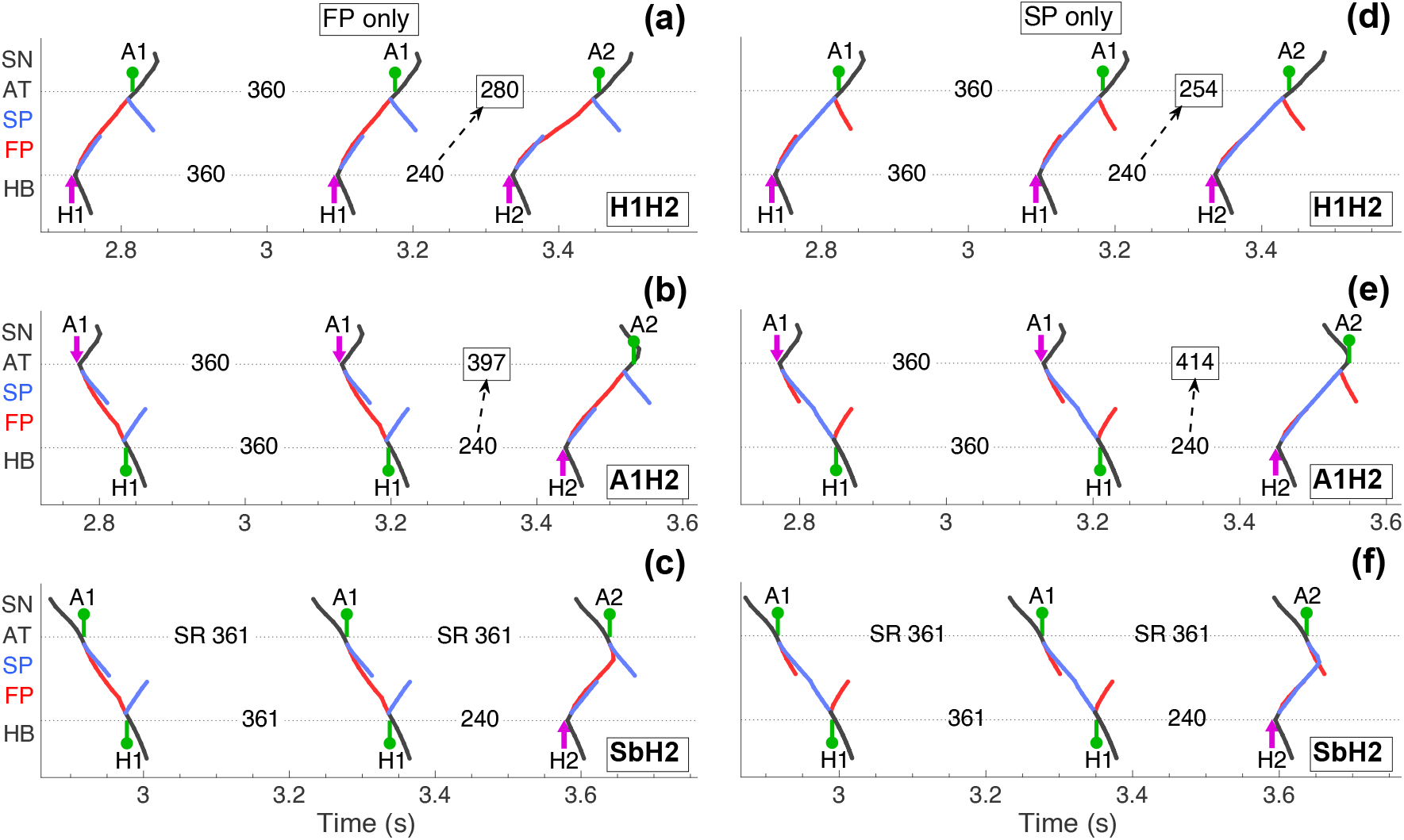
Ladder diagrams for the first model variant during His bundle pacing with *γ* = 1.7 and a 240 ms test interval. Panels (a)–(c) show activation patterns with only the fast pathway (FP) present and panels (d)–(f) with only the slow pathway (SP). The top row illustrates standard H1H2 pacing, the middle row shows A1H2 pacing, and the bottom row shows SbH2 pacing. Dashed arrows highlight the transformation of the applied H1–H2 test interval into the corresponding A1–A2 interval.

The SbH2 method introduces an additional challenge due to potential collision between anterograde sinus beats and retrograde H2-evoked excitation, which complicates measurement of H2-A2 conduction times and ERPs. This issue can be mitigated by temporarily suppressing the sinus beat following the final A1 event.

#### 3.2.2. Retrograde ERPs and Reentry Induction Across Model Variants

Retrograde ERP curves as functions of the autonomic nervous system coefficient *γ*, and reentrant activity maps for all three model variants are presented in Fig. 7. For comparison, shaded regions in the upper panels denote AVNRT or echo-beat induction obtained using standard H1H2 pacing, while those in the lower panels correspond to the alternative A1H2/SbH2 protocols. Across all variants, the alternative pacing methods consistently yield shorter FP and SP ERPs (Table 1) and predict smaller reentry susceptibility regions, which shift toward increasing parasympathetic tone.

**Table 1:**
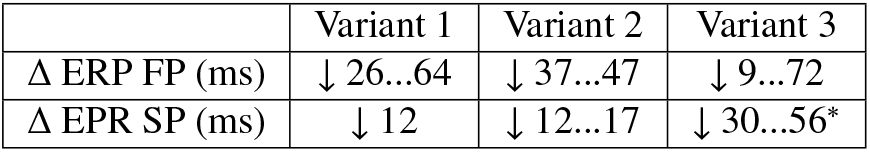
Shortening of the effective refractory periods (ERP) of fast (FP) and slow (SP) pathways for the three model variants as identified with A1H2 pacing. (*) Not accounting for upward shift considered in Appendix.

**Figure 7:**
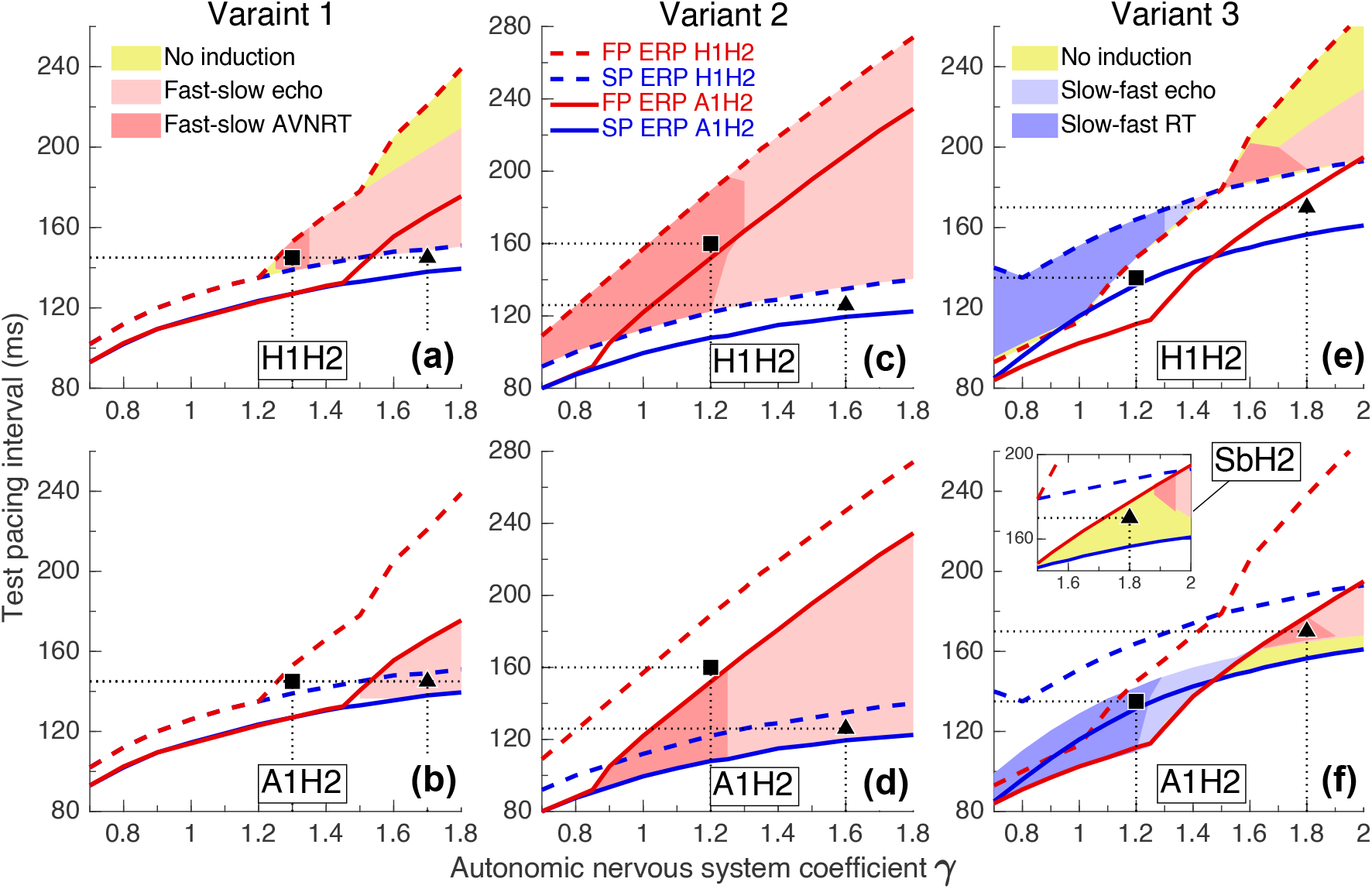
Retrograde effective refractory periods (ERPs) as functions of the autonomic nervous system coefficient *γ*, and reentrant activity maps for the first (a,b), second (c,d), and third (e,f) model variants. Dashed lines show ERPs obtained with standard H1H2 pacing and solid lines show ERPs obtained with A1H2 or SbH2 pacing. Red and blue lines represent the fast (FP) and slow (SP) pathways. Light and dark shading indicates regions where fast–slow echo beats or AVNRT were induced. In panels (b) and (d), A1H2 and SbH2 results coincide. In panel (f), the main plot shows A1H2 results and the inset illustrates the divergence seen with SbH2 pacing for *γ* ≥ 1.5. Markers denote parameter points analyzed in Figs. 8–10.

Fast–slow (atypical) AVNRT appears only when FP and SP ERPs are close in value, at small test intervals and enhanced sympathetic tone. In contrast to atrial pacing, the regions between ERP curves that fail to produce either AVNRT or echo beats are significantly reduced or even eliminated under His-bundle pacing.

#### 3.2.3. Conduction Patterns in the Unmodified AV Node

Ladder diagrams illustrating the differences between pacing methods for all three model variants appear in Figs. 8–10. In the first model variant, a test condition that induces fast–slow AVNRT under H1H2 pacing yields only normal retrograde conduction under A1H2/SbH2 pacing. Conversely, conditions that appear non-inducible using H1H2 pacing can produce echo beats when alternative methods are applied. The second model variant exhibits similar behavior, though with more extensive echo-beat activity at some test points.

**Figure 8:**
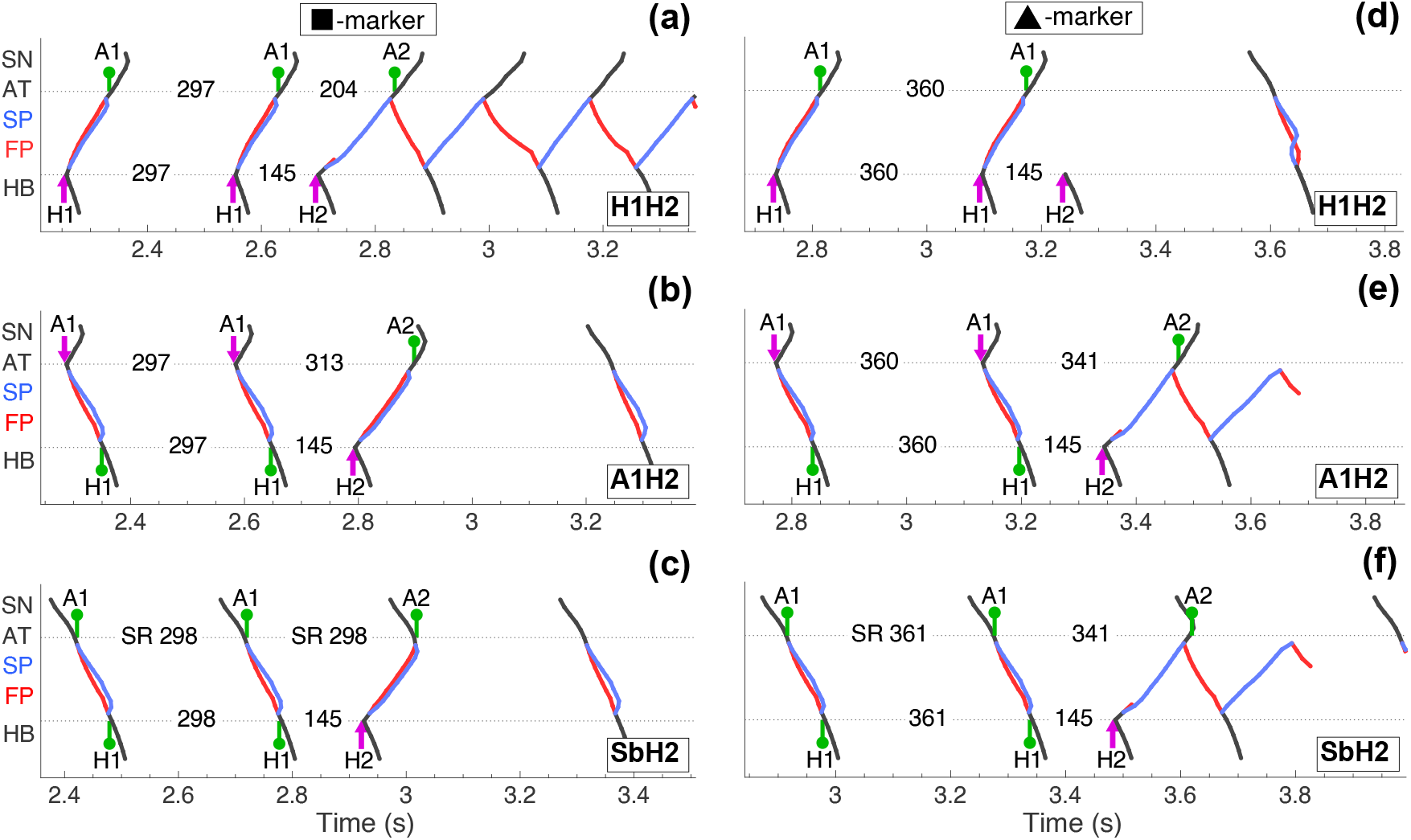
Ladder diagrams of the unmodified AV node for the first model variant obtained with H1H2, A1H2, and SbH2 pacing. Panels (a)–(c) correspond to autonomic coefficient *γ* = 1.3 with a 145 ms H1–H2 interval, matching the square markers in Fig. 7(a, b). Panels (d)–(f) show results for *γ* = 1.7 with the same test interval, corresponding to the triangle markers in Fig. 7(a, b).

**Figure 9:**
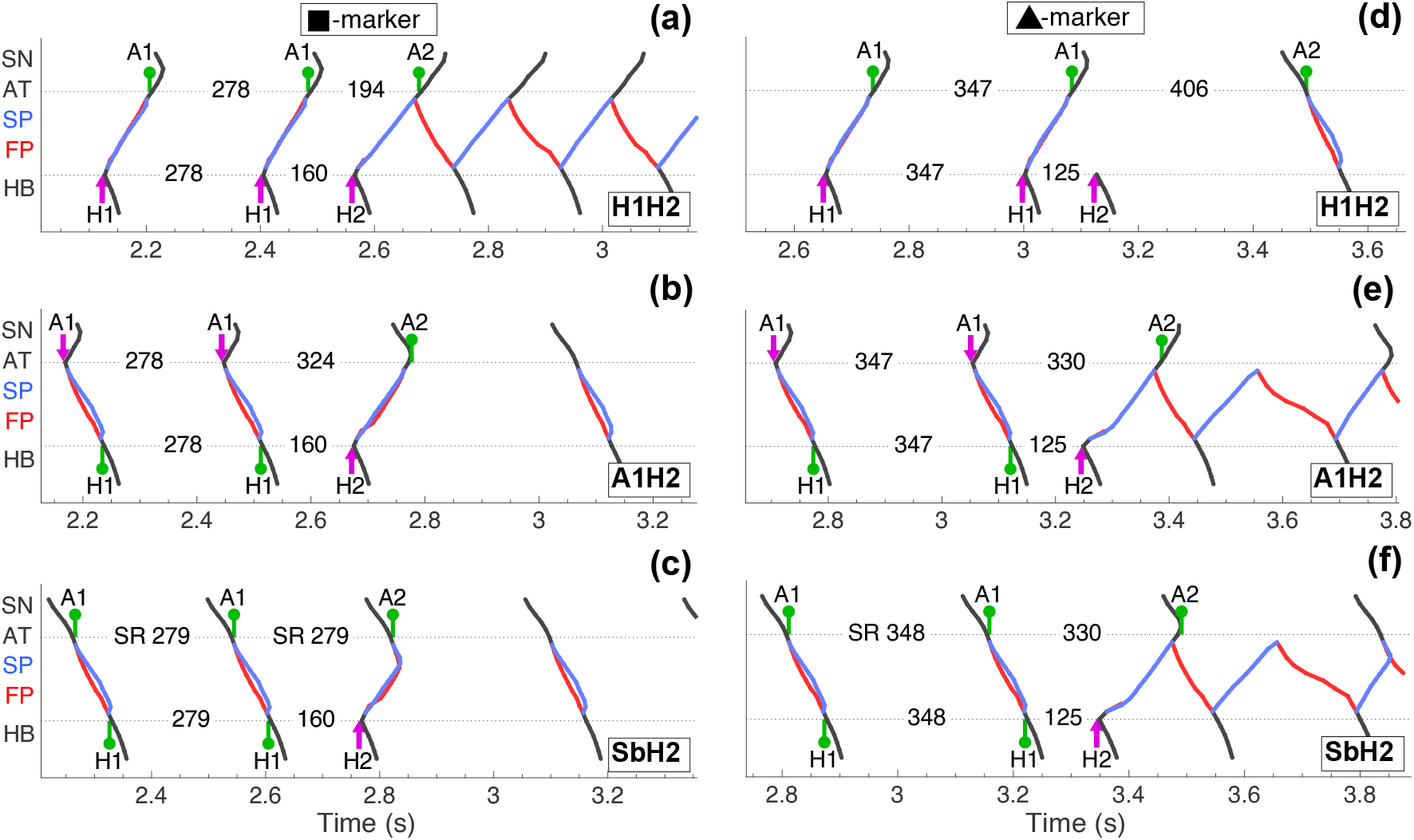
Ladder diagrams of the unmodified AV node for the second model variant obtained with H1H2, A1H2, and SbH2 pacing. Panels (a)–(c) correspond to autonomic coefficient *γ* = 1.2 with a 160 ms H1–H2 interval, matching the square markers in Fig. 7(c, d). Panels (d)–(f) show results for *γ* = 1.6 with with a 125 ms H1–H2 interval, corresponding to the triangle markers in Fig. 7(c, d).

**Figure 10:**
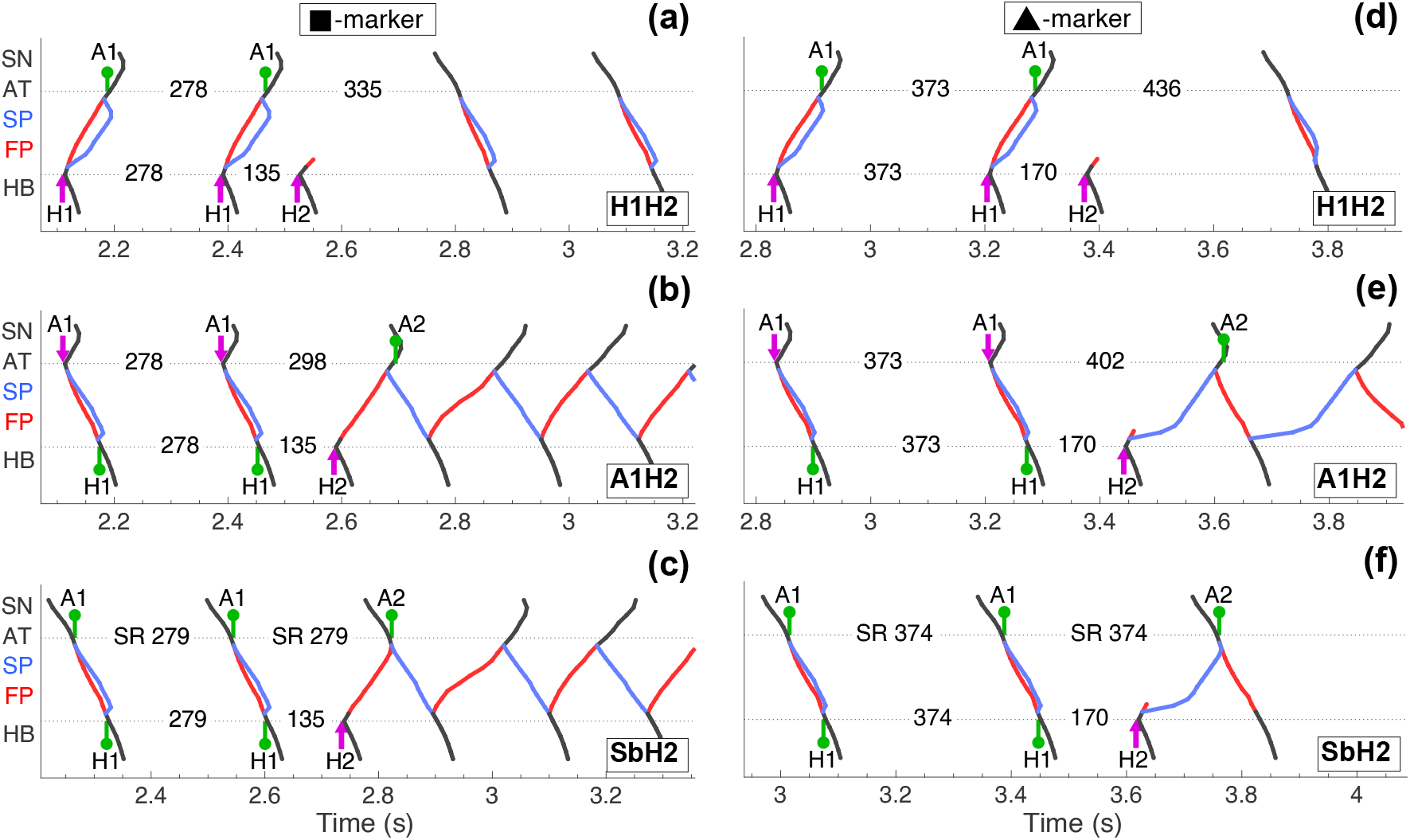
Ladder diagrams of the unmodified AV node for the third model variant obtained with H1H2, A1H2, and SbH2 pacing. Panels (a)–(c) correspond to autonomic coefficient *γ* = 1.2 with a 135 ms H1–H2 interval, matching the square markers in Fig. 7(e, f). Panels (d)–(f) show results for *γ* = 1.8 with with a 170 ms H1–H2 interval, corresponding to the triangle markers in Fig. 7(e, f).

The third model variant displays the most complex behavior. Under H1H2 pacing, some test conditions appear non-inducible, yet the alternative methods reveal both slow–fast and fast–slow AVNRT. In SbH2 pacing, anterograde sinus activation may collide with retrograde echo beats, which prevents propagation within SP and can effectively suppress fast–slow reentry. This interaction highlights the influence of the sinus node in shaping retrograde conduction outcomes under mixed-direction pacing.

#### 3.2.4. Retrograde Conduction Curves

Figure 11 compares retrograde conduction curves obtained using H1H2 pacing, A1H2 pacing with sinus activity temporarily blocked, and A1H2/SbH2 pacing with an active sinus node. Because collisions between anterograde and retrograde waves may prevent complete curve reconstruction under A1H2 or SbH2 pacing, sinus node suppression is sometimes necessary to obtain full curves comparable to those from H1H2 pacing. Across all autonomic states, conduction curves obtained with the A1H2 method consistently show shorter ERPs and faster retrograde conduction than those produced by standard H1H2 pacing.

**Figure 11:**
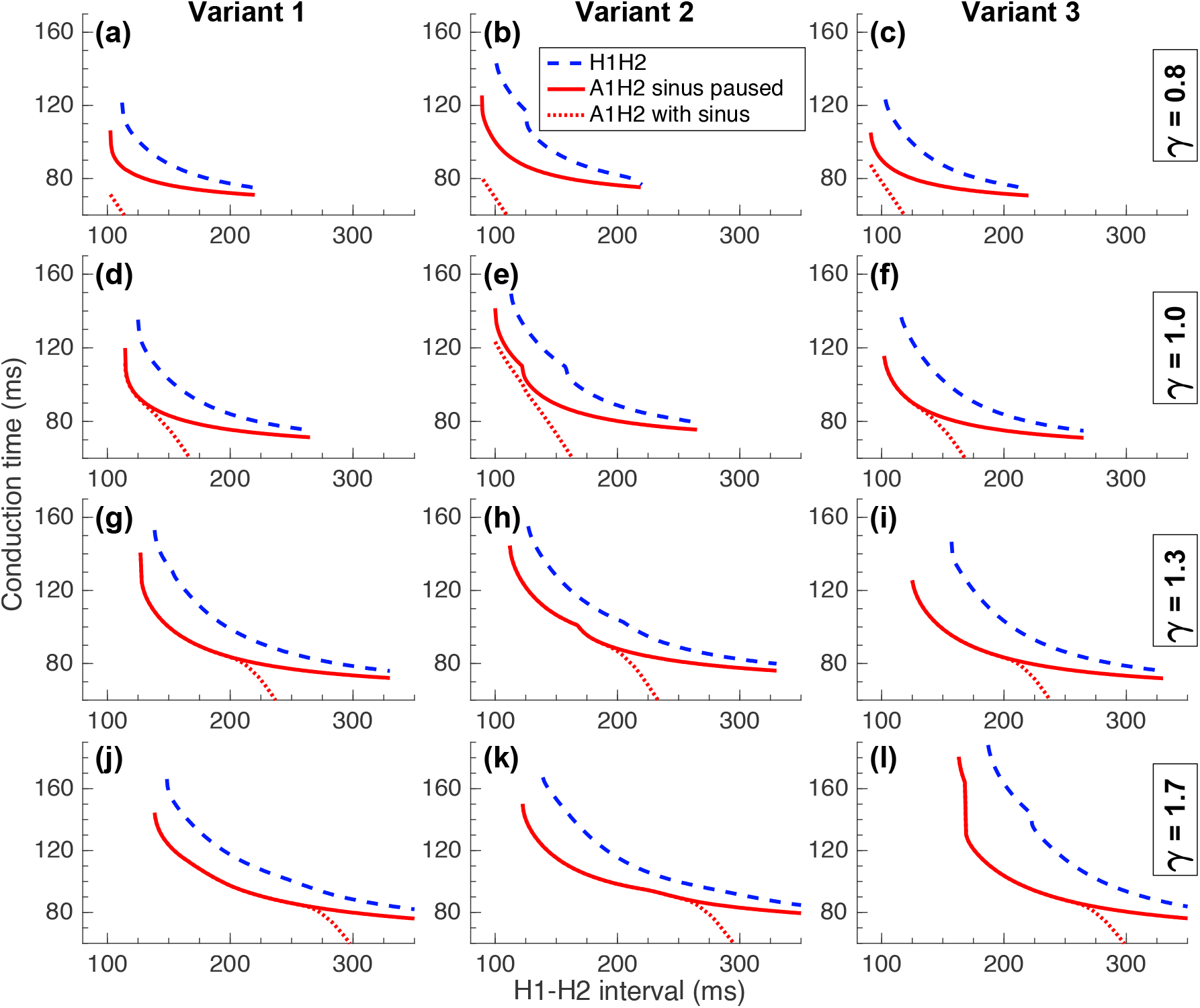
Retrograde conduction times in the unmodified AV node during H1H2 pacing (blue dashed lines), A1H2 pacing with sinus temporarily paused (red solid lines), and SbH2 pacing (red dotted lines) for autonomic nervous system states defined by *γ* = 0.8 (panels a–c), 1.0 (d–f), 1.3 (g–i), and 1.7 (j–l).

#### 3.2.5. Effectiveness of the Alternative Pacing Methods

Figure 12 illustrates the maps of reentrant activity, comprising both echo beats and areas of atrioventricular nodal reentrant tachycardia (AVNRT), produced by standard H1H2 pacing, highlighting regions of overestimation and underestimation. Table 2 presents the corresponding quantitative results for the areas of reentrant activity *A* and the associated errors Σ for the standard H1H2 method compared to alternative pacing methods, as calculated using Eqs. 3-5. In all three model variants, the area overestimated is larger than the area underestimated. Furthermore, the overlap area, where the reentrant activity areas derived from both the standard and alternative methods coincide, is largest in the second model variant, suggesting that the effectiveness of the alternative pacing methods improves as the reentrant activity areas become smaller.

**Table 2:** Areas of reentrant activity *A* and errors of the standard H1H2 pacing method Σ relative to A1H2 and SbH2 (in brackets) methods.

|  | Variant 1 | Variant 2 | Variant 3 |
| --- | --- | --- | --- |
| $A_{H1H2}$ (a.u.) | 19.2 | 80.3 | 28.9 |
| $A_{A1H2}$ (a.u.) | 7.2 | 55.4 | 23.8 (20.7) |
| $A_{Overlap}$ (a.u.) | 3.7 | 42.4 | 4.9 |
| $\Sigma_{H1H2}^{Over}$ (%) | 68.6 | 40.6 | 50.3 (53.8) |
| $\Sigma_{H1H2}^{Under}$ (%) | 15.3 | 14.0 | 39.6 (35.3) |
| $\Sigma_{H1H2}^{Total}$ (%) | 83.9 | 54.6 | 89.9 (89.1) |

**Figure 12:**
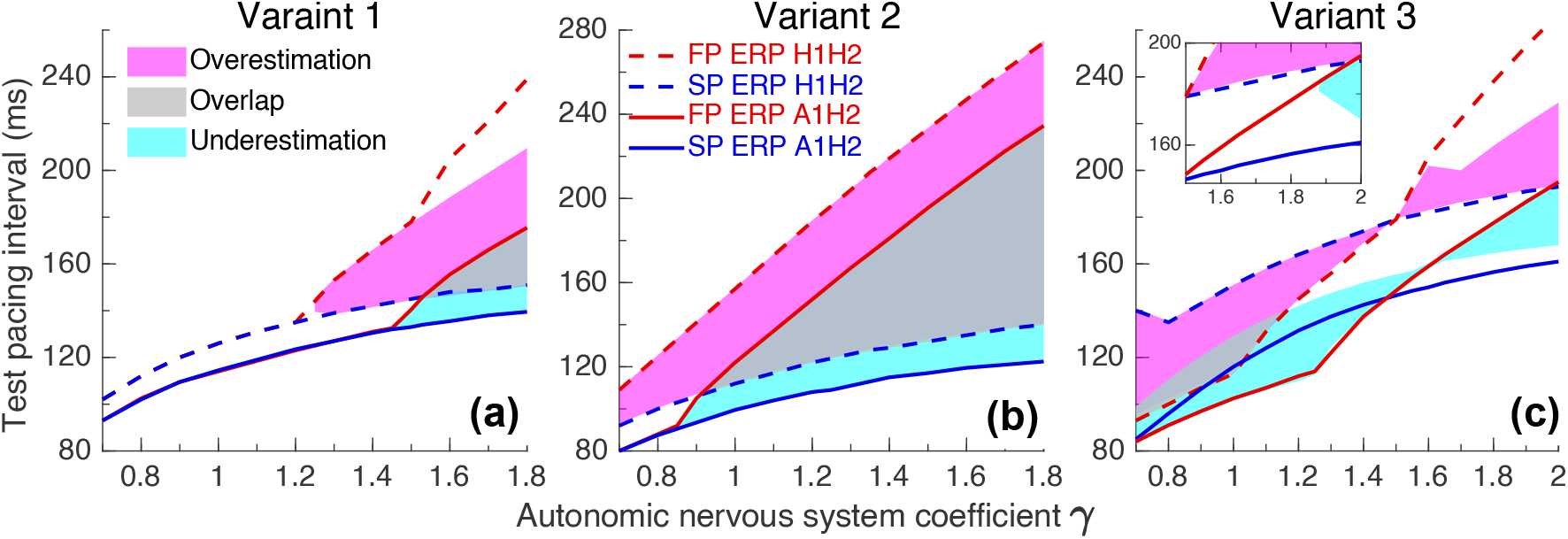
Over- and underestimation maps of reentrant activity produced by standard H1H2 pacing, as identified with A1H2 pacing. In panel (c), the main plot shows results for A1H2 and the inset illustrates the divergence seen with SbH2 pacing for *γ* ≥ 1.5.

## 4. Discussion

In this study, we introduce alternative pacing strategies that more accurately characterize AV nodal conduction and the conditions that promote atrioventricular nodal reentrant tachycardia (AVNRT). By comparing these methods with the standard S1S2 protocol for atrial (A1A2) and His bundle (H1H2) stimulation, we demonstrate that the inferred refractory properties of the fast (FP) and slow (SP) pathways, as well as the predicted likelihood of reentry, are highly dependent on the pacing approach.

Previous work [11] assumed that any premature stimulus delivered within the classical induction window defined [12, 13] by the effective refractory periods (ERPs) of the pathways would initiate reentry. However, simulations using the standard S1S2 protocol revealed substantial discrepancies between expected and observed regions of AVNRT and echo beats. The pacing schemes proposed here, including SbA2 for atrial stimulation and A1H2/SbH2 for His bundle stimulation, allow premature testing in the presence of ongoing anterograde activation. This provides a more physiologically realistic assessment of nodal function and reveals conduction behaviors that remain hidden under the S1S2 approach.

Across the full range of autonomic nervous system states, three reproducible regions emerged between the FP and SP retrograde ERP curves: AVNRT induction, echo beats, and non-inducible responses. These reflect distinct dynamical regimes of the nodal reentrant loop, ranging from sustained oscillations to critically damped or fully blocked responses. Importantly, the transitions between these regimes are not determined solely by retrograde ERPs. Concealed anterograde FP activation [17, 22] plays a critical role by suppressing retrograde conduction at specific coupling intervals. This mechanism explains the appearance of non-inducible regions within the traditional induction window and highlights the need to account for bidirectional interactions during refractoriness testing [5].

Atrial pacing results obtained with A1A2 and SbA2 protocols were nearly identical, with only minor differences related to the origin of the coupling current in the proximal atrial inputs. In contrast, His bundle pacing demonstrated pronounced protocol dependence. The alternative pacing schemes consistently produced shorter retrograde ERPs and smaller regions of AVNRT and echo beats compared with the standard H1H2 protocol, indicating that S1S2-based His stimulation may either exaggerate or underestimate the vulnerability to reentry. In several configurations, persistent sinus activation even suppressed reentry entirely, underscoring the importance of pacing direction and ongoing activation in shaping retrograde conduction outcomes.

An additional phenomenon emerged in the third model variant, where parasympathetic enhancement inverted the FP–SP ERP relationship. Under these conditions, slow-fast AVNRT and echo beats occurred in regions where both pathways appeared capable of retrograde conduction. This paradox arose from the interplay between SP conduction observed in the FP-ablated preparation used for ERP measurement and retrograde block in the unmodified node at specific test intervals. This behavior, elaborated in the Appendix, illustrates the sensitivity of nodal dynamics to both structural modifications and the measurement protocol itself.

Finally, the SbA2 and SbH2 pacing strategies, which maintain sinus rhythm, proved valuable for reproducing more natural patterns of activation within the nodal ring. These approaches captured conduction phenomena that the S1S2 protocol obscures, reinforcing the importance of physiological pacing context when evaluating AV nodal recovery and reentry. Overall, our findings demonstrate that alternative pacing methods provide a more reliable and physiologically grounded framework for estimating AVNRT inducibility, particularly during His bundle stimulation. These results have implications for computational modeling and may inform clinical electrophysiology, where accurate assessment of nodal refractoriness and conduction is essential for diagnosing and managing reentrant tachyarrhythmias.

## 5. Limitations

This study relies on a computational model constructed from experimental data obtained in rabbit hearts [15, 25]. While the fundamental principles of dual-pathway conduction are conserved across species, anatomical and physiological differences [36], such as AV nodal pacemaker location, atrial–nodal connectivity, and intrinsic cycle length, may influence the quantitative details of conduction and refractoriness in humans. Although these differences are unlikely to alter the qualitative mechanisms described here, direct extrapolation to the human AV node should be made with caution.

The pacing strategies examined in this work were evaluated solely through simulations and have not yet been validated experimentally. Their practical implementation, feasibility in an electrophysiology laboratory, and agreement with *in vivo* measurements remain to be determined. Additionally, the model does not account for certain structural and functional complexities of the AV node, including microscopic heterogeneity, transitional cell populations, three-dimensional geometry, and spatially distributed autonomic input [7]. Finally, a full sensitivity analysis of parameter dependence was not performed and may provide further insight into the robustness of the observed behaviors.

## 6. Conclusion

We developed and evaluated alternative atrial and His-bundle pacing methods designed to more accurately assess AV nodal refractoriness and reentrant susceptibility, including AVNRT and echo beats. Through comprehensive simulations across a broad range of autonomic states, we found that the classical S1S2 method provides reliable results for atrial pacing but may distort the assessment of retrograde conduction during His bundle stimulation. In particular, S1S2 His pacing can either overestimate or underestimate the true regions of AVNRT and echo beat induction. The proposed pacing strategies incorporate sinus beats to preserve physiologic anterograde activation, thereby enabling more accurate measurement of FP and SP refractory periods and the retrograde interactions that shape reentry induction. Compared with the S1S2 method, the alternative approaches yield smaller and more precisely localized induction regions and shift the predicted thresholds toward shorter test intervals, better reflecting the underlying nodal dynamics.

Taken together, these findings provide a refined conceptual and methodological framework for evaluating AV nodal conduction and reentry. The proposed pacing methods may improve the interpretation of electrophysiological testing with His bundle stimulation and provide a foundation for future experimental studies aimed at validating and translating these insights into clinical practice.

## CRediT authorship contribution statement

**Maxim Ryzhii:** Writing – original draft, Writing – review & editing, Visualization, Validation, Software, Investigation, Data curation, Supervision. **Elena Ryzhii:** Writing – original draft, Writing – review & editing, Methodology, Investigation, Conceptualization, Formal analysis.

